# Molecular Glue-Induced Homodimerization Drives Targeted CRBN Autodegradation

**DOI:** 10.64898/2026.03.08.710235

**Authors:** Lu Chen, Xiaoxiao Zou, Jiayin Liang, Jun Wang, Xiaolin Luo, Taiting Shi, Xinzhu Li, Shuke Yang, Linhui Cao, Yingjie Sun, Yue Zhao, Hewei Jiang, Yi Jiang, Zhipeng Su, Huan Xiong, Cheng Luo, Wenchao Lu

## Abstract

Molecular glue degraders (MGDs) offer a sophisticated, proximity-based approach to protein modulation. In this study, we introduce LJY-3-60, a novel proximity-inducing agent that unexpectedly triggers the potent and selective autodegradation of CRBN. Evidence from CRISPR-Cas9 screening and IP-MS reveals that this degradation process is strictly governed by the intrinsic CRL4^CRBN^ machinery, independent of any extrinsic E3 recruitment. Through a combination of cellular and biophysical characterizations, we demonstrate that LJY-3-60 acts as a molecular bridge to template CRBN homodimerization. This mechanism is unequivocally elucidated by the atomic-resolution co-crystal structure of the CRBN^Midi^-LJY-3-60 complex. The structure explicitly delineates the homodimerization interface, revealing how the ligand reorganizes the protein surface to stabilize a non-canonical architecture that drives trans-autoubiquitination and subsequent proteasomal degradation. Furthermore, LJY-3-60 serves as a highly effective, controllable off-switch to mitigate PROTAC-induced toxicity. Ultimately, this work delivers a robust chemical tool for modulating CRBN stability. By demonstrating how a small molecule can functionally mimic an endogenous E3 substrate’s degron to catalyse targeted autodegradation, this study establishes a rational structural framework for designing the next generation of self-destructive modulators in targeted protein degradation (TPD) therapeutics.

## Introduction

The paradigm of targeted protein degradation (TPD), championed by PROTACs and molecular glues, has fundamentally reshaped the drug discovery landscape by bringing “undruggable” proteins within therapeutic reach(1-3). Among the handful of E3 ligases currently harnessed for TPD, Cereblon (CRBN)—the primary target of immunomodulatory imide drugs (IMiDs)—stands as the most extensively characterized and utilized(4). Historically, CRBN based degraders have operated by orchestrating *de novo* protein-protein interactions (PPIs) to redirect neo-substrates toward ubiquitination and subsequent proteasomal destruction(5-8). While the field has been intensely focused on expanding the repertoire of targetable neo-substrates such as IKZFs, CK1a, Wee1, NEK7, PDE6D and among others(9-11), the chemical space of molecular glues occasionally yields unexpected pharmacological behaviors(12). For instance, the unintended depletion of CRBN has sometimes been observed as an off-target artifact, or intentionally achieved through the use of bulky, bivalent Homo-PROTACs. However, there are sparse reports of monovalent CRBN-based molecular glues capable of selectively orchestrating CRBN homodimerization to trigger its autodegradation. Such a molecule would not only reveal a fascinating new dimension of proximity-induced pharmacology but also provide a highly desirable pharmacological tool to acutely modulate E3 ligase activity, offering a precise means to temporally control TPD interventions.

During our ongoing exploration of novel molecular glues, we serendipitously identified LJY-3-60, a potent and selective monovalent compound that triggers rapid CRBN autodegradation. Our CRISPR-Cas9 genetic screening and IP-MS profiling reveal that LJY-3-60 bypasses the need for extrinsic E3 recruitment; instead, it elegantly co-opts the intrinsic CRL4^CRBN^ machinery. By engaging UBE2D2—a primary E2 enzyme that acts in concert with CRL4 complexes—LJY-3-60 facilitates the ligase’s self-destruction(13).

Crucially, we unequivocally elucidate this atypical mechanism by solving the atomic-resolution co-crystal structure of the CRBN^Midi^-LJY-3-60 complex(5, 6, 14). Our structural analysis reveals that LJY-3-60 functions as a molecular bridge, templating a non-canonical CRBN homodimerization interface. This induced proximity drives efficient trans-autoubiquitination within the dimerized assembly, culminating in rapid proteasomal degradation. Through systematic structure-activity relationship (SAR) studies, we further refined the essential pharmacophore of this self-destructive framework. Ultimately, this work not only delivers a robust chemical off-switch for controlling CRBN stability but also provides a transformative structural blueprint for how small molecules can functionally mimic E3 degrons to catalyze targeted autodegradation(15).

## Results

### Discovery of LJY-3-60: A Highly Potent CRBN Ligand Identified via Orthogonal Binding Assays

During our efforts to develop novel molecular glues, we implemented an orthogonal screening strategy utilizing both biochemical and cellular assays. This workflow aimed to identify scaffolds with high CRBN target occupancy by cross-validating hits through time-resolved fluorescence energy transfer (TR-FRET) and cell-based NanoBRET assays (**Figure 1A**). From this library screen, LJY-3-60 emerged as a standout candidate, exhibiting exceptionally high binding affinity towards CRBN that warranted further investigation. In the TR-FRET binary binding assay, LJY-3-60 effectively competed for the CRBN binding site with an IC_50_ of 0.047 μM, demonstrating almost 30-fold greater potency than the benchmark, lenalidomide (IC_50_ = 1.6 μM) (**Figure 1B**). To validate these *in vitro* biochemical findings within a physiological context, we assessed target occupancy in live cells using the NanoBRET assay. These experiments confirmed that LJY-3-60 maintains its high-potency profile intracellularly (IC_50_ = 0.086 μM) (**Figure 1C**). The striking consistency across these orthogonal assays, coupled with its exceptionally robust binding affinity, prompted us to investigate whether LJY-3-60 modulates CRBN protein homeostasis or drives neosubstrate recruitment beyond mere pocket occupancy.To comprehensively evaluate the biological consequences of LJY-3-60 treatment, we performed unbiased quantitative proteomics to assess its global impact on the cellular proteome. Strikingly, CRBN emerged as the sole protein significantly downregulated across the entire proteome, while other E3 ligases and potential off-targets remained stable (**Figure 1D**). This exceptional specificity underscored the highly selective nature of LJY-3-60-induced CRBN depletion. Western blot analysis corroborated this degradative profile, revealing a clear dose-dependent depletion of CRBN with remarkably rapid kinetics. Substantial reduction in CRBN protein levels was observed as early as 2 hours post-treatment, highlighting the compound’s robust efficiency (**Figure 1E**). To delineate the mechanistic pathway driving this degradation, we employed pharmacological rescue experiments. CRBN degradation was completely abrogated by pretreatment with pomalidomide (Pom, a CRBN competitor), MLN4924 (a NEDD8-activating enzyme inhibitor), or bortezomib (a proteasome inhibitor). These results unambiguously confirm that LJY-3-60-induced CRBN depletion is strictly dependent on target occupancy, cullin-RING ligase (CRL) neddylation, and proteasomal activity, perfectly aligning with a canonical ubiquitin-proteasome system (UPS) mechanism (**Figure 1E**).

**Figure 1.**
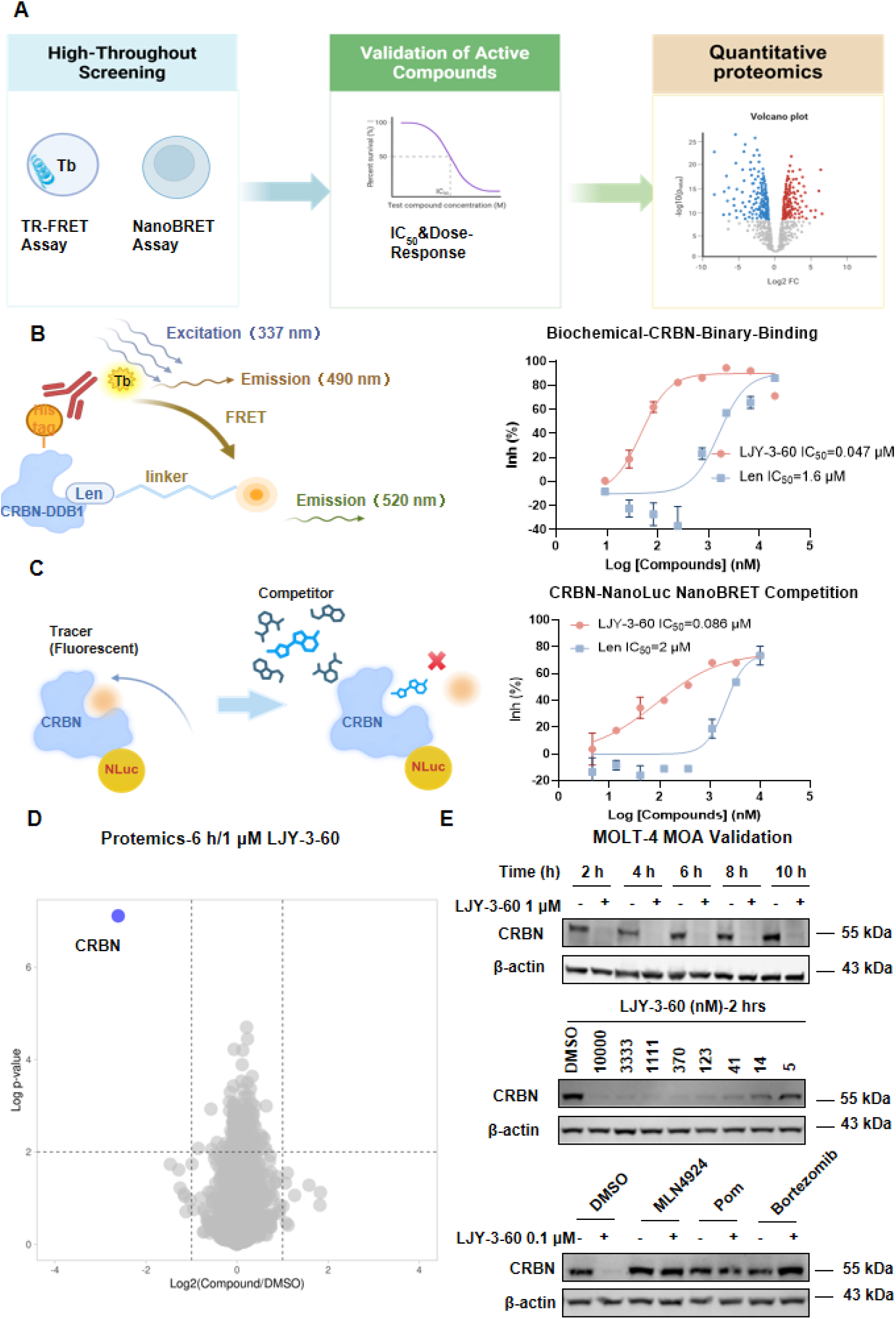
Discovery of LJY-3-60 as a potent and selective inducer of CRBN autodegradation. **A**, Schematic workflow of the study, from high-throughput screening to quantitative proteomics. **B**, Schematic representation of the TR-FRET assay (left) and binary binding curves (right) for LJY-3-60 and lenalidomide. Data represent the mean ± SD of two independent experiments (n = 2). **C**, Schematic of the NanoBRET competition assay (left) and corresponding intracellular target engagement analysis in HEK293T cells (right). LJY-3-60 demonstrates superior target engagement compared to lenalidomide (Len). Data represent the mean ± SD (n = 2). **D**. Volcano plot of global quantitative proteomics in MOLT-4 cells treated with LJY-3-60 (1 μM) for 6 h with three biological replicates, highlighting the highly selective depletion of CRBN. **E**, Immunoblot analysis demonstrating the rapid (within 2 h) and dose-dependent degradation of CRBN. This degradation is completely abrogated by pretreatment with MLN4924, lenalidomide, or bortezomib, confirming a UPS-dependent and on-target mechanism.

### Elucidation of the LJY-3-60-Induced CRBN Homodimerization and Autodegradation Mechanism

To genetically delineate the mechanism driving this atypical autodegradation, we performed a genome-wide CRISPR-Cas9 positive selection screen utilizing an eGFP-CRBN reporter system **(Figure 2A**). Interestingly, the screen predominantly enriched UBE2D2—a primary E2 ubiquitin-conjugating enzyme known to pair with CRL4 complexes—while lacking significant enrichment for any extrinsic E3 ligases (**Figure 2B**). This specific genetic dependency profile, centered on CRBN’s native E2 partner, led us to hypothesize that LJY-3-60 might co-opt CRBN’s intrinsic catalytic machinery rather than recruiting a trans-acting E3 ligase. Consequently, we proposed a homodimerization-driven model, wherein the compound acts as a molecular template to assemble a CRBN-CRBN complex, thereby facilitating trans-autoubiquitination.

**Figure 2.**
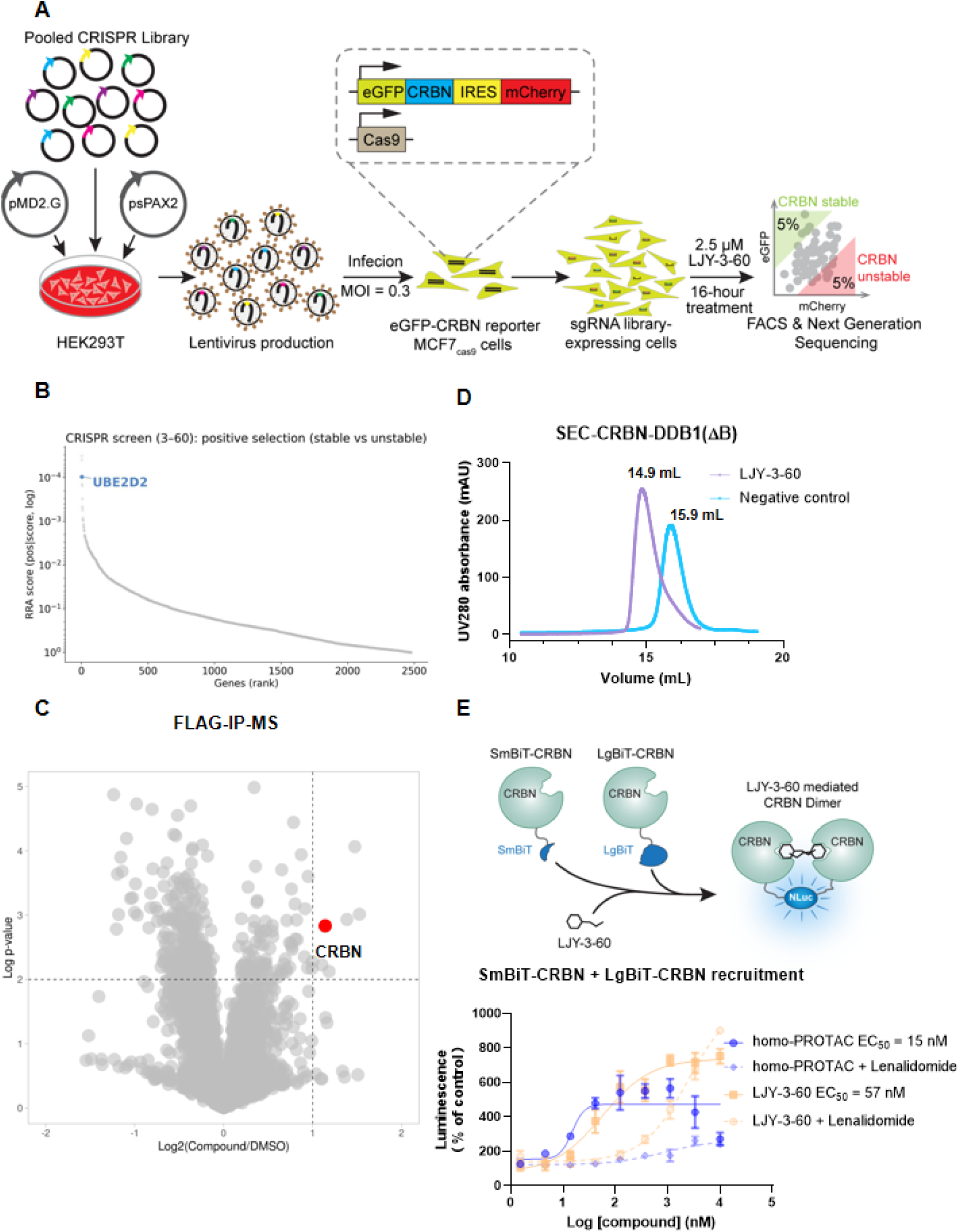
LJY-3-60 triggers targeted CRBN autodegradation via homodimerization. **A**, Schematic representation of the genome-wide CRISPR-Cas9 positive selection screen. **B**, Volcano plot from the CRISPR screen identifying UBE2D2 as a critical genetic dependency for LJY-3-60-induced degradation. **C**, FLAG-IP-MS interactome analysis (or Volcano plot of FLAG-IP-MS) demonstrating the significant and selective enrichment of CRBN, corroborating the homodimerization model. **D**, Size-exclusion chromatography (SEC) profiles of the CRBN-DDB1 complex. The distinct peak shift from 15.9 to 14.9 mL upon LJY-3-60 treatment confirms stable dimer formation in vitro. **E**, Schematic of the NanoBiT complementation assay (up) and the dose-response curve (down) validating LJY-3-60-mediated CRBN-CRBN assembly in live cells.

This hypothesis was rigorously substantiated through orthogonal biochemical and biophysical approaches. Flag-IP-MS analysis revealed that CRBN was the exclusively enriched E3 ligase following LJY-3-60 treatment (**Figure 2C**). Furthermore, in vitro size-exclusion chromatography (SEC) unambiguously demonstrated a compound-dependent peak shift, indicative of the formation of a stable, dimeric assembly comprising two CRBN-DDB1(ΔB) complexes (**Figure 2D**). Finally, live-cell NanoBiT assays corroborated the rapid formation of this homodimeric complex in cellulo. Crucially, this signal was effectively antagonized by the addition of excess lenalidomide, confirming that the assembly is strictly dependent on the occupancy of the canonical immunomodulatory drug (IMiD)-binding pocket (**Figure 2E**).

### LJY-3-60 Induces CRBN Homodimerization by Mimicking a Neosubstrate G-Loop Degron

To elucidate the molecular mechanism governing LJY-3-60-induced CRBN autodegradation, we first assessed its capacity to drive the in vitro homodimerization of CRBN^Midi^, an engineered construct optimized for recombinant expression in *E. coli*. Size-exclusion chromatography (SEC) revealed that the addition of LJY-3-60 induces a pronounced shift of the CRBN^Midi^ peak toward a smaller elution volume, confirming the compound’s ability to mediate the formation of a stable homodimeric complex (**Figure 3A**)(16, 17). To delineate the atomic-level basis of this induced proximity, we solved the co-crystal structure of the CRBN^Midi^-LJY-3-60 complex with a resolution of 1.8 Å (**Figure 3B, Table S1**). The structure reveals a C2-symmetric homodimer wherein two molecules of LJY-3-60 (designated MGD/A and MGD/B) act as molecular bridges connecting the two CRBN subunits. Unambiguous electron density for both ligand copies is evident in the Polder omit map. The two molecules are further stabilized by inter-ligand π-π stacking of their aromatic ring systems at the core of the dimer interface, significantly bolstering the overall stability of the dimeric assembly (**Figure 3B-C**). Detailed structural analysis reveals that the dimer interface is orchestrated by a highly cooperative *cis-trans* interaction network (**Figure 3C**). On the *cis-side*, the glutarimide core of each LJY-3-60 molecule is deeply sequestered within the canonical tri-tryptophan (tri-Trp) pocket (W380/388/400), establishing three conserved hydrogen bonds with W380 and H378. Additionally, the compound scaffold forms a hydrogen bond with N351 and engages in extensive hydrophobic and π-π stacking interactions with surrounding residues, including P352, H353, and Y355. Crucially, the two ligand copies play distinct, non-equivalent roles in “stitching” the dimer interface: the O_28_ atom of MGD/A forms an essential hydrogen bond with the side chain of H397’ on the *trans-subunit*, whereas the O_30’_ carbonyl of MGD/B establishes a vital hydrogen bond with the W400 side chain on the *cis-subunit*. The indispensability of these specific interfacial contacts was confirmed using negative control analogs LJY-4-154 and LJY-5-6 (**Table 1**).

**Figure 3.**
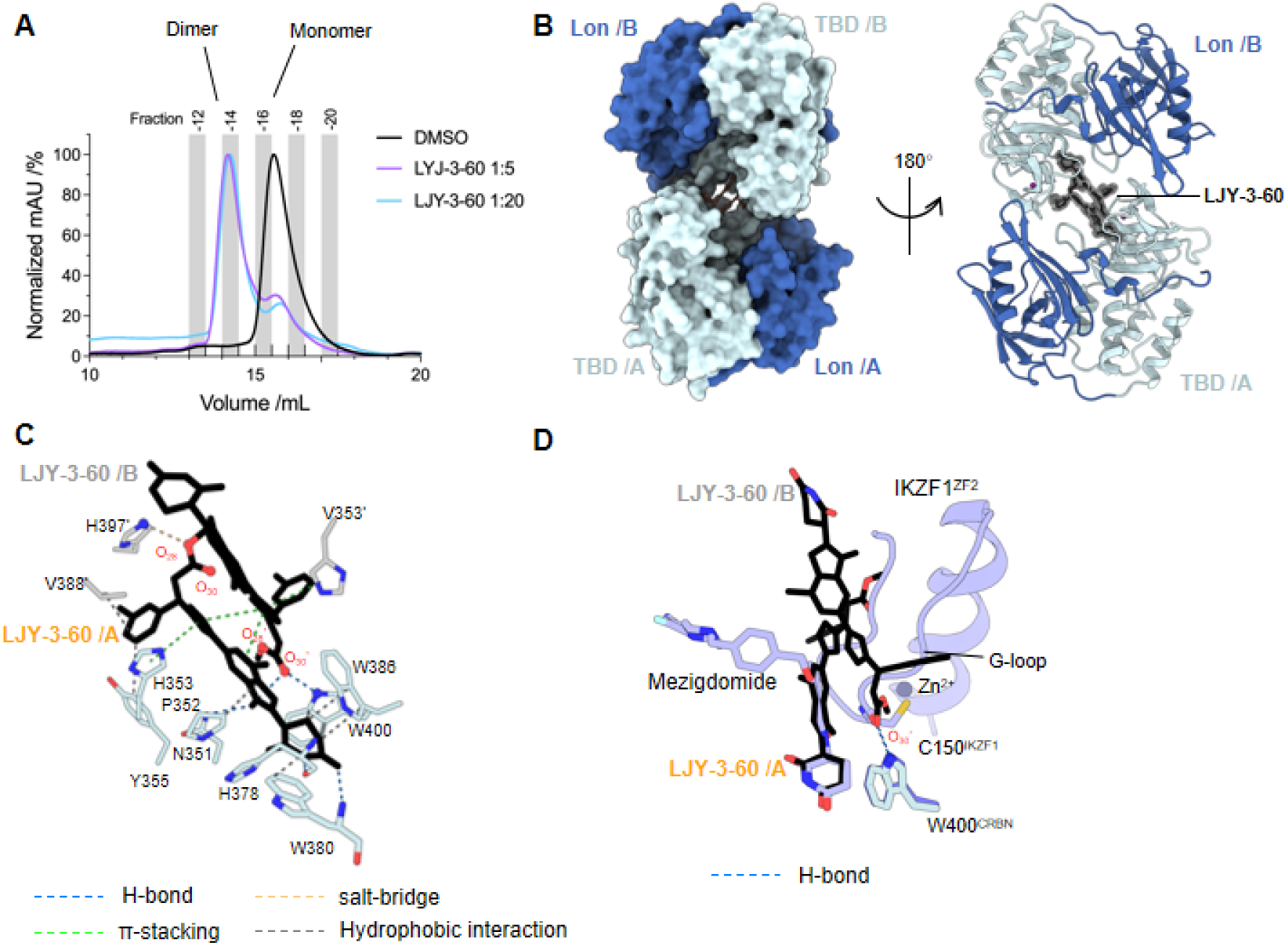
Structural basis of LJY-3-60-induced CRBN homodimerization. **A**, Size-exclusion chromatography profiles of CRBNMidi protein incubated with DMSO (black) or LJY-3-60 at varying molar ratios. **B**, Overall architecture of the LJY-3-60-mediated CRBN^Midi^ homodimer (PDB ID: 22UL, this paper). Left: Surface representation of the homodimer, indicating protomer A (/A) and protomer B (/B). The Lon (Blue) and thalidomide-binding domain (TBD, cyan) are labeled. Right: Ribbon representation of the complex. The polder mFo - Fc omit map for LJY-3-60 (stick, orange) is shown as a gray mesh and contoured at 3.0σ. **C**, Detailed interactions within the CRBN tri-tryptophan pocket. LJY-3-60 (orange/gray sticks) interacts with key residues of CRBN^Midi^. **D**, Superimposition of the LJY-3-60-CRBN^Midi^ structure (orange/gray/cyan) with the CRBN^Midi^-Mezigdomide-IKZF1 complex (PDB ID: 8RQC, purple). LJY-3-60 occupies a similar spatial position to the G-loop of IKZF1^ZF2^, highlighting its role as a structural mimic that facilitates dimerizati.

Despite retaining the CRBN-binding glutarimide core, these analogs failed to induce dimerization, underscoring their inability to establish the productive interface interactions observed with LJY-3-60. Remarkably, the structural data unequivocally elucidate that LJY-3-60 functions as a bona fide degron mimic. Structural superposition of our CRBN^Midi^-LJY-3-60 complex with the reported CRBN^Midi^-mezigdomide-IKZF1 ternary complex (PDB ID: 8RQC) reveals that the spatial trajectory of LJY-3-60 closely aligns with the G-loop degron of the IKZF1 neosubstrate (**Figure 3D**). Specifically, through its critical hydrogen bond with W400 on the *cis-side*, MGD/B meticulously recapitulates both the spatial orientation and the physicochemical properties of natural G-loop recognition. This exquisite structural mimicry effectively disguises one CRBN protomer as a pseudo-neosubstrate for the other, orchestrating the formation of a productive ternary assembly that triggers trans-autoubiquitination and subsequent self-degradation.

### Structure-Activity Relationship (SAR) and Robustness of the LJY-3-60 Framework

Guided by the atomic-resolution insights from the co-crystal structure, we conducted a systematic structural modification campaign to delineate the pharmacophoric elements driving CRBN autodegradation. Biochemical evaluation revealed that all 22 synthesized analogs maintained potent, concentration-dependent CRBN-DDB1 binding affinities when tested at 0.1, 1.0, and 10 μM (**Table 1**). However, translating this target occupancy into productive degradation was exquisitely dependent on specific structural features. Chief among these was stereochemistry. Within the Scaffold 1 series, the (R)-enantiomer LJY-3-60 (**Table 1, entry 1**) exhibited potent CRBN depletion capacity (achieving 91% degradation at 1 μM), whereas its (S)-counterpart LJY-4-154 (**Table 1, entry 2**) was completely inactive. This stringent stereochemical requirement was faithfully mirrored in the Scaffold 2 series, as exemplified by the active (R)-isomer LXL-2-65 (**Table 1, entry 17**) and the inactive (S)-isomer LJY-4-161 (**Table 1, entry 5**).

Next, we probed the necessity of the aryl halogens. Removal of the chlorine atom from LJY-3-60 yielded LJY-4-153 (**Table 1, entry 3**), which retained equipotent degradation efficacy (**Figure 4A-B**). This indicated that the chlorine substitution is dispensable for productive ternary complex formation, a finding that guided the design of the chlorine-free Scaffold 2 series (**Table 1, entries 3-21**). Furthermore, stripping the bromine atom from LJY-4-153 generated LXL-2-65 (**Table 1, entry 17**), which still maintained robust degradation activity (achieving ∼90% degradation).

**Figure 4.**
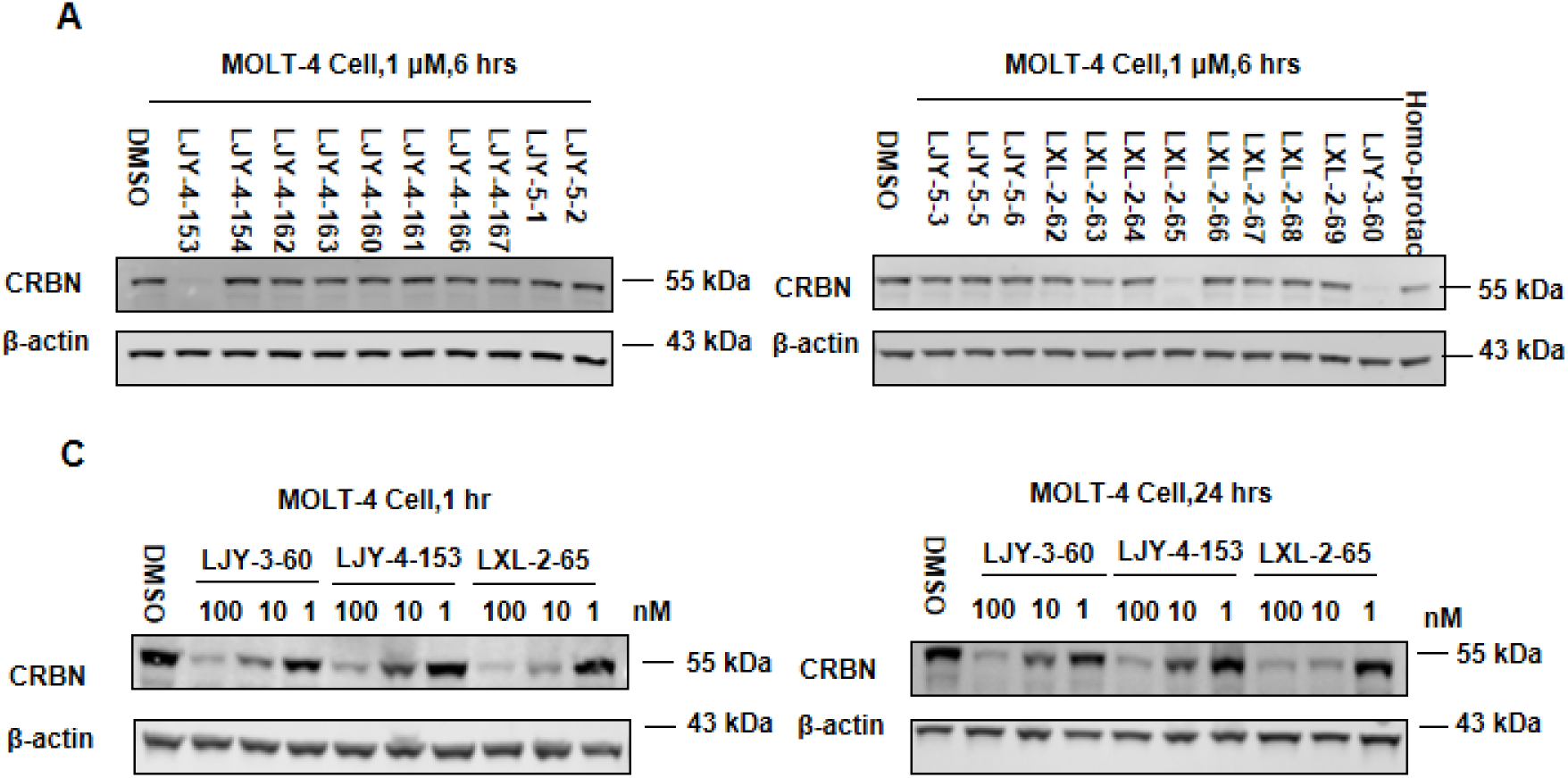
SAR analysis and structural robustness of the CRBN degraders. **A**, Immunoblot analysis of endogenous CRBN levels in MOLT-4 cells following treatment with key scaffold analogs (1 μM) for 6 h. **B**, Immunoblot analyses demonstrating the dose and time-dependent kinetics of CRBN depletion across the scaffold series. The data confirm consistent and potent degradation efficiency at nanomolar concentrations, even following scaffold dehalogenation.

In stark contrast, key structural alterations to the core functional groups of LXL-2-65 resulted in a complete abrogation of degradative activity. Hydrolysis of the ester, bioisosteric replacement of the ester with an amide, introduction of a hydroxyl group, incorporation of an aliphatic ring, or modulation of the aliphatic chain length universally abolished activity **(Table 1**). Additionally, replacing the central triazole ring with an aliphatic linker and introducing an R3 group (yielding LJY-5-5, **Table 1, entry 22**) similarly yielded an inactive compound. Overall, these SAR trends firmly establish that while the precise (R)-stereochemistry and the core triazole-ester scaffold are absolutely paramount for templating CRBN homodimerization, the peripheral halogens are dispensable. In perfect concordance with our structural data, the dimeric interface is highly tolerant of dehalogenation, as the loss of these atoms does not disrupt the essential cis-trans interaction network. Ultimately, these findings highlight the remarkable structural robustness of the LJY-3-60 framework, providing a rational trajectory for developing next-generation self-destructive molecular glues with minimized molecular weights and optimized pharmacokinetic profiles.

### Pharmacological Application: LJY-3-60 as a Universal Off-Switch for TPD Therapeutics

Building on our structural insights, we explored the translational utility of this self-destructive mechanism as a programmable off-switch for TPD therapeutics. Remarkably, LJY-3-60 successfully rescued target protein levels 32 hours following the washout of a long-acting PROTAC (**Figure 5A-B**). The superiority of this ligase-depletion strategy over traditional competitive inhibition became particularly evident when challenged with the ultra-potent molecular glue, CC-885. We observed that CC-885-mediated GSPT1 degradation was completely refractory to rescue by the canonical CRBN binder pomalidomide, likely due to the exceptionally high affinity and cooperative stabilization of the CC-885-CRBN-GSPT1 ternary complex.

**Figure 5.**
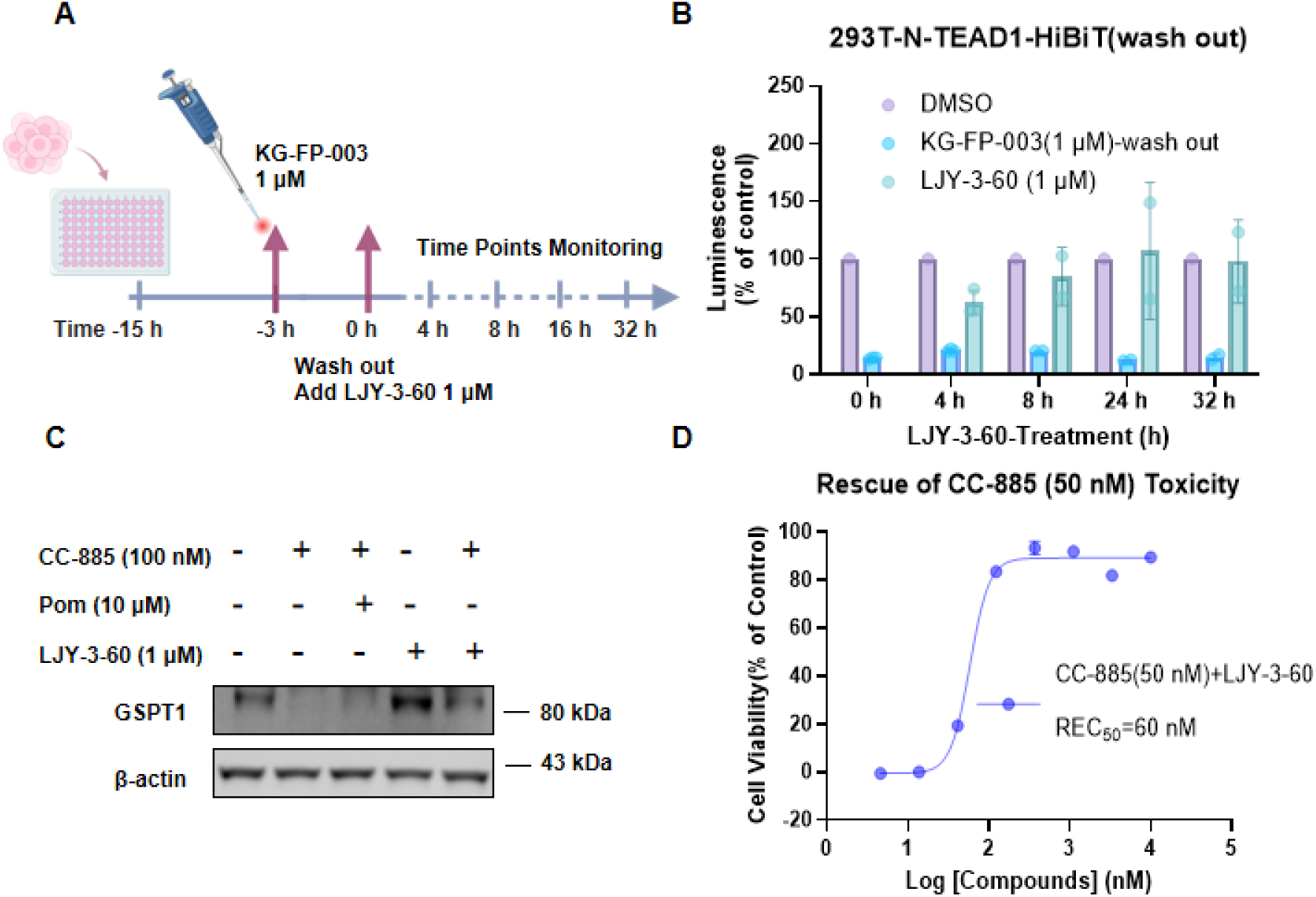
LJY-3-60 functions as a universal safety switch for TPD therapeutics. **A**, Schematic workflow of the PROTAC washout and rescue assay. **B**, Rescue of TEAD1 protein levels following PROTAC washout. HEK293T cells expressing N-TEAD1-HiBiT were pre-treated with the TEAD PROTAC KG-FP-003 (1 μM) for 3 h. Subsequent treatment with LJY-3-60 (1 μM) effectively restored TEAD1 levels. **C**, Immunoblot analysis demonstrating that LJY-3-60 (1 μM), but not pomalidomide (10 μM), successfully rescues GSPT1 from degradation induced by the ultra-potent molecular glue CC-885 (100 nM). **D**, Dose-response curve highlighting the potent reversal of CC-885-induced cytotoxicity by LJY-3-60, yielding a calculated half-maximal rescue concentration (REC_50_) of 60 nM. Quantitative data are normalized to the DMSO control and presented as mean ± SD (n = 3).

In stark contrast, LJY-3-60 effectively eradicated the cellular CRBN pool, thereby halting GSPT1 degradation and fully restoring cell viability with a half-maximal rescue concentration REC_50_ of 60 nM (**Figure 5C-D**). Together, these findings establish LJY-3-60 as a potent, universal antidote, offering a transformative pharmacological strategy to enhance the safety, reversibility, and precise temporal control of TPD-based therapies.

## Discussion

In summary, our work establishes LJY-3-60 as a first-in-class molecular glue that drives the self-degradation of the E3 ligase CRBN through a unique homodimerization mechanism. By integrating genome-wide screening, proteomic profiling, and atomic-resolution crystallography, we mapped a molecular blueprint in which a small molecule acts as a structural template to remodel the E3 surface, initiating trans-autoubiquitination independently of extrinsic E3 recruitment.

This finding challenges the conventional binary view of E3 ligase ligands as merely simple inhibitors or substrate recruiters. LJY-3-60 demonstrates that E3 ligases can be reprogrammed to target themselves for destruction. This “degrader-of-the-degrader” paradigm introduces an innovative pharmacological strategy for achieving precise temporal control over targeted protein degradation (TPD) processes. Experimentally, LJY-3-60 effectively restored GSPT1 protein levels and neutralized CC-885–induced cytotoxicity with a half-maximal rescue concentration (REC_50_) of 60 nM, validating its utility as a rapid and efficient chemical off-switch to halt overactive degradation.

Beyond its application as a regulatory switch, LJY-3-60 serves as a high-resolution chemical probe to investigate the acute cellular consequences of CRBN loss, bypassing the limitations and compensatory effects of genetic knockouts. Building on the structural insights from the 1.8 Å CRBN^Midi^-LJY-3-60 complex, we have established a rational design framework that can be applied to engineer self-degrading molecules against other key E3 ligases by mimicking degrons from endogeous subrates. Ultimately, these chemical tools will deepen our understanding of E3 ligase homeostasis and expand the conceptual repertoire for designing next-generation, dynamically controllable TPD technologies.

## Acknowledgement

This work was supported by the Strategic Priority Research Program of the Chinese Academy of Sciences (Grant Nos. XDB0830301 and XDB1260302 to C.L.), the Science and Technology Commission of Shanghai Municipality (Grant No. YDZX20233100004032 to C.L.), and the National Natural Science Foundation of China (Grant No. 82304301 to H.X.). W.L. acknowledges support from the Shanghai Pujiang Talent Program (Grant No. 23PJ1430100) and internal funding from Lingang Laboratory. We thank the staff members of the National Facility for Protein Science in Shanghai (NFPS) and the Shanghai Synchrotron Radiation Facility (SSRF) for their technical support and assistance in data collection and analysis. Specifically, we are grateful for the assistance provided by the staff at beamline BL10U2.

