## Supplemental Information for "Molecular Glue-Induced Homodimerization Drives Targeted CRBN Autodegradation"

[2. Table S1. Data collection and refinement statistics for crystal structure of CRBN](#_Toc16729)^[Midi](#_Toc16729)^ [complex with LJY-3-60	7](#_Toc16729)

### 1. Supplementary Figures

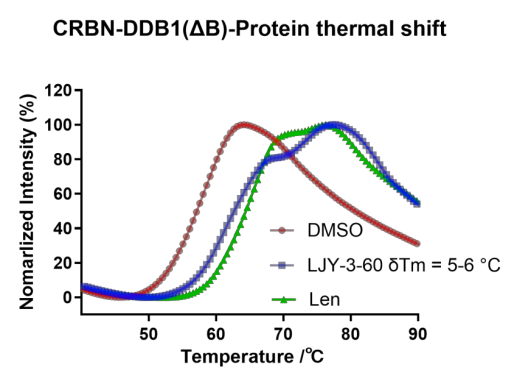

Supplementary Fig. 1. Treatment with LJY-3-60 resulted in a substantial rightward shift in the melting curve (Δ Tm = 5–6 °C), confirming higher physical binding affinity for the E3 ligase complex relative to Lenalidomide.

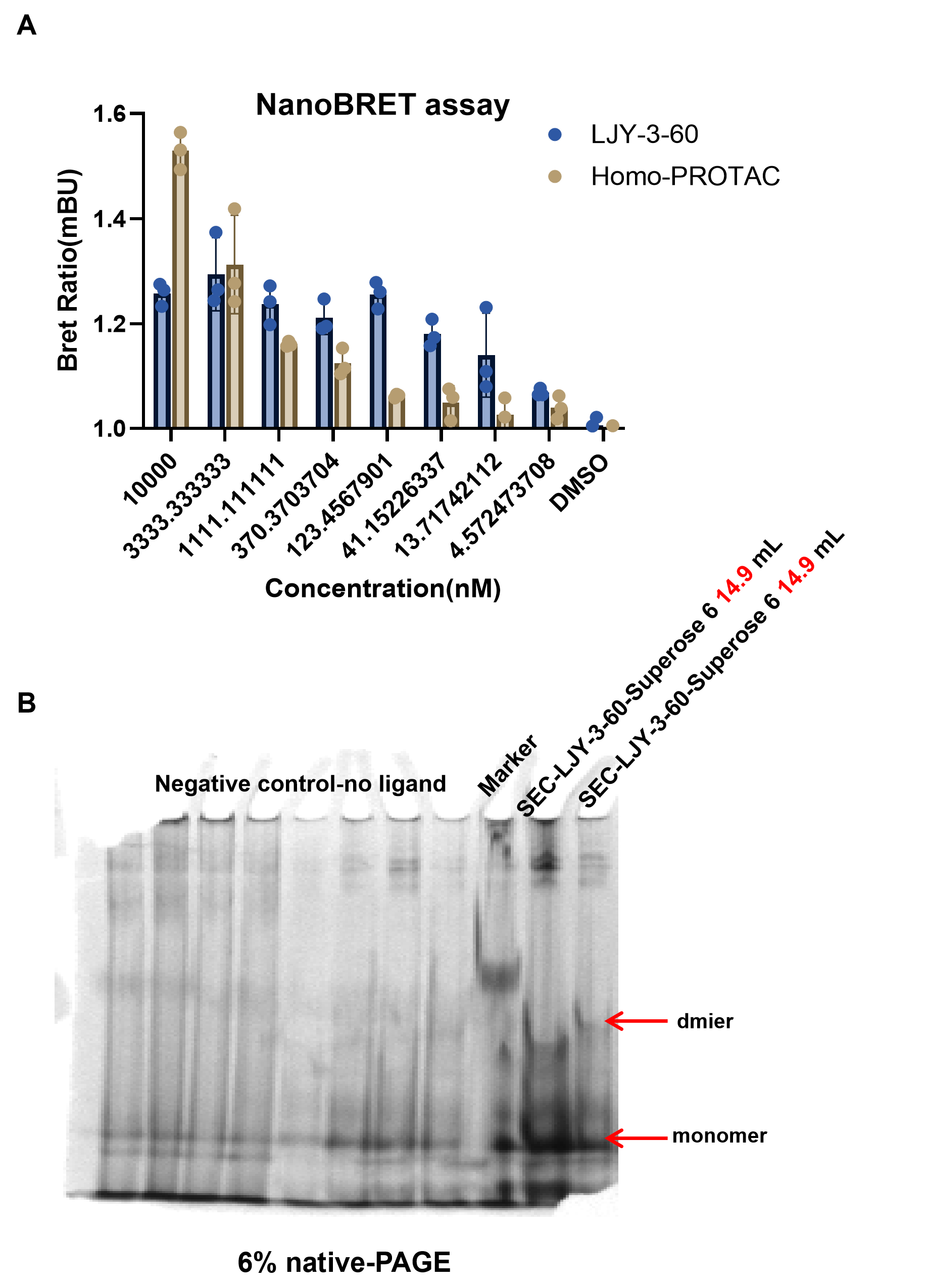

Supplementary Fig. 2**.** (A) Dose-dependent NanoBRET assay in HEK293T cells co-expressing NanoLuc–CRBN and HaloTag–CRBN. Cells were treated with LJY-3-60 or a reference Homo-PROTAC (4.5–10,000 nM). Data represent mean ± SD (n = 3 biological replicates). (B) 6% native-PAGE of purified CRBN–DDB1 (ΔB) protein. Treatment with LJY-3-60 (SEC-purified samples from Superose 6 or Superdex 200) induced a distinct band shift corresponding to the CRBN–CRBN dimer, absent in the negative control (no ligand).

Supplementary Fig. 3**.** Uncropped immunoblots

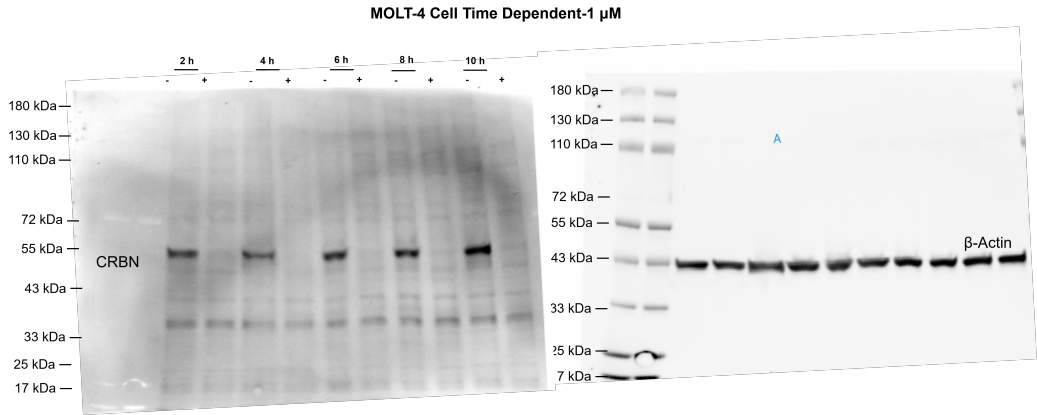

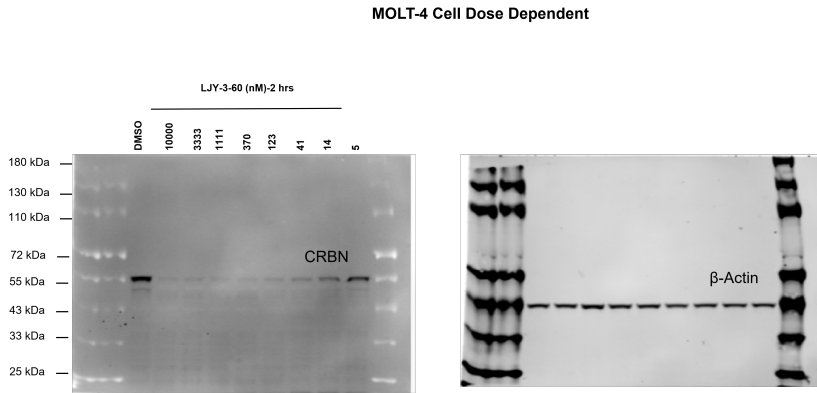

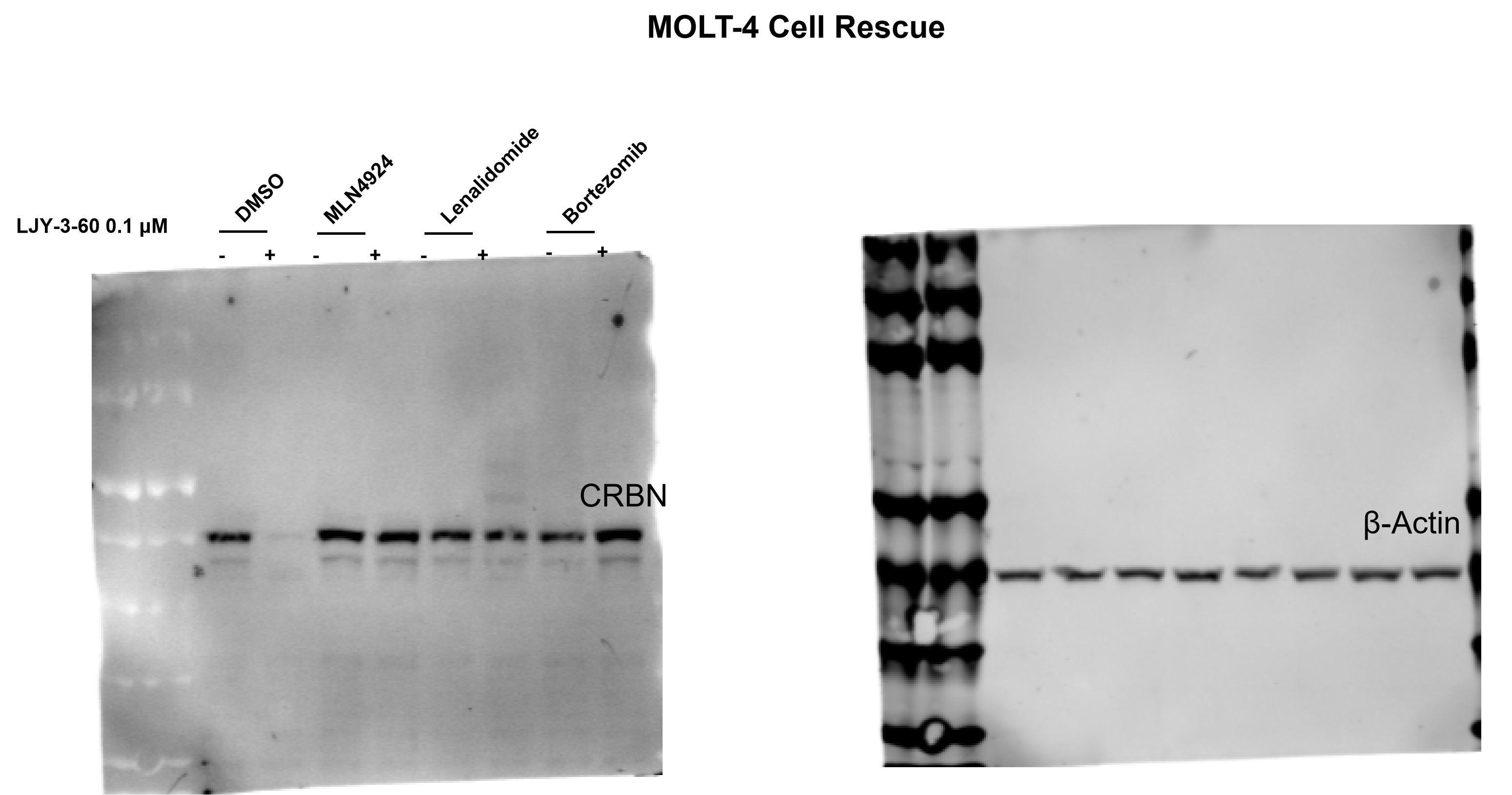

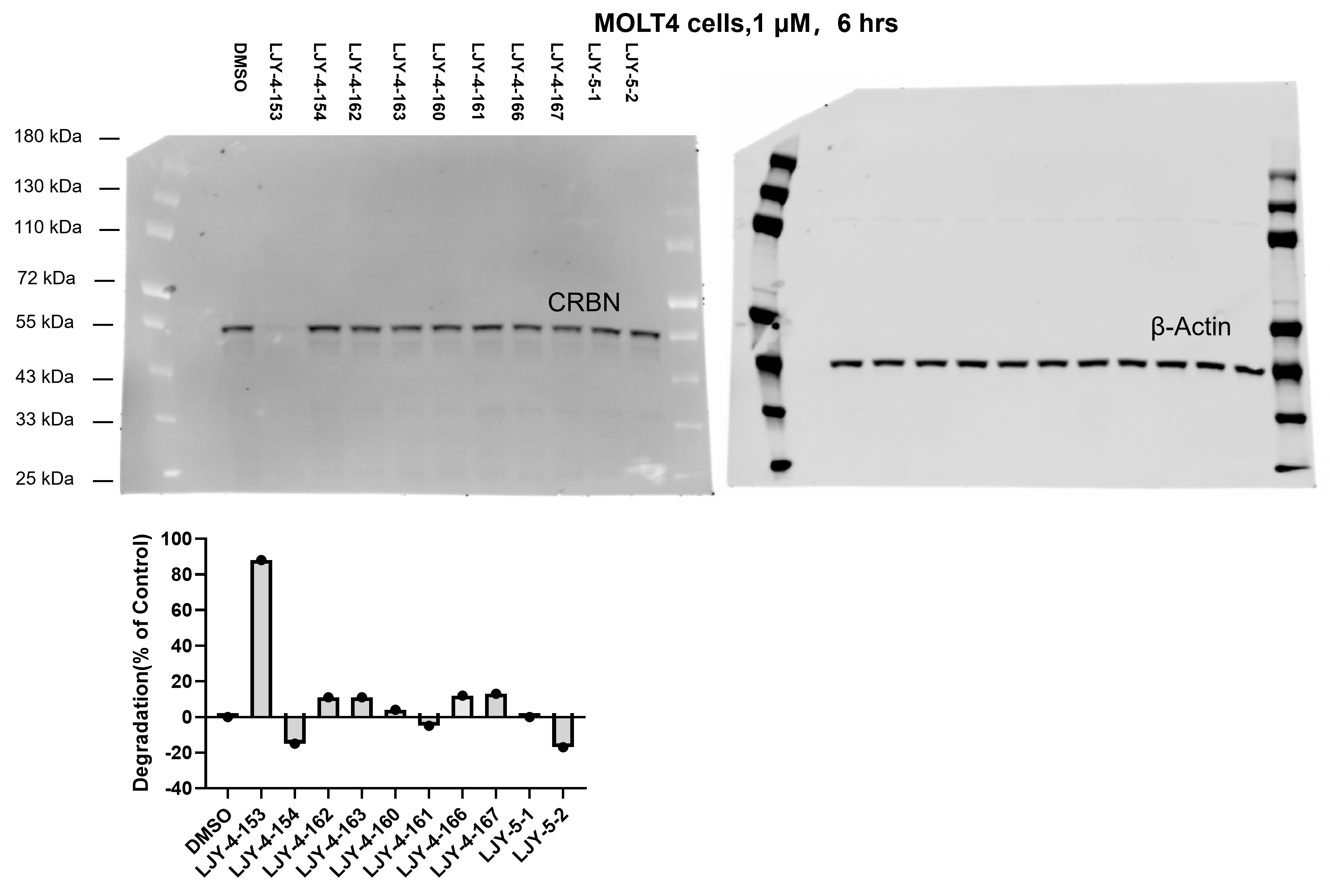

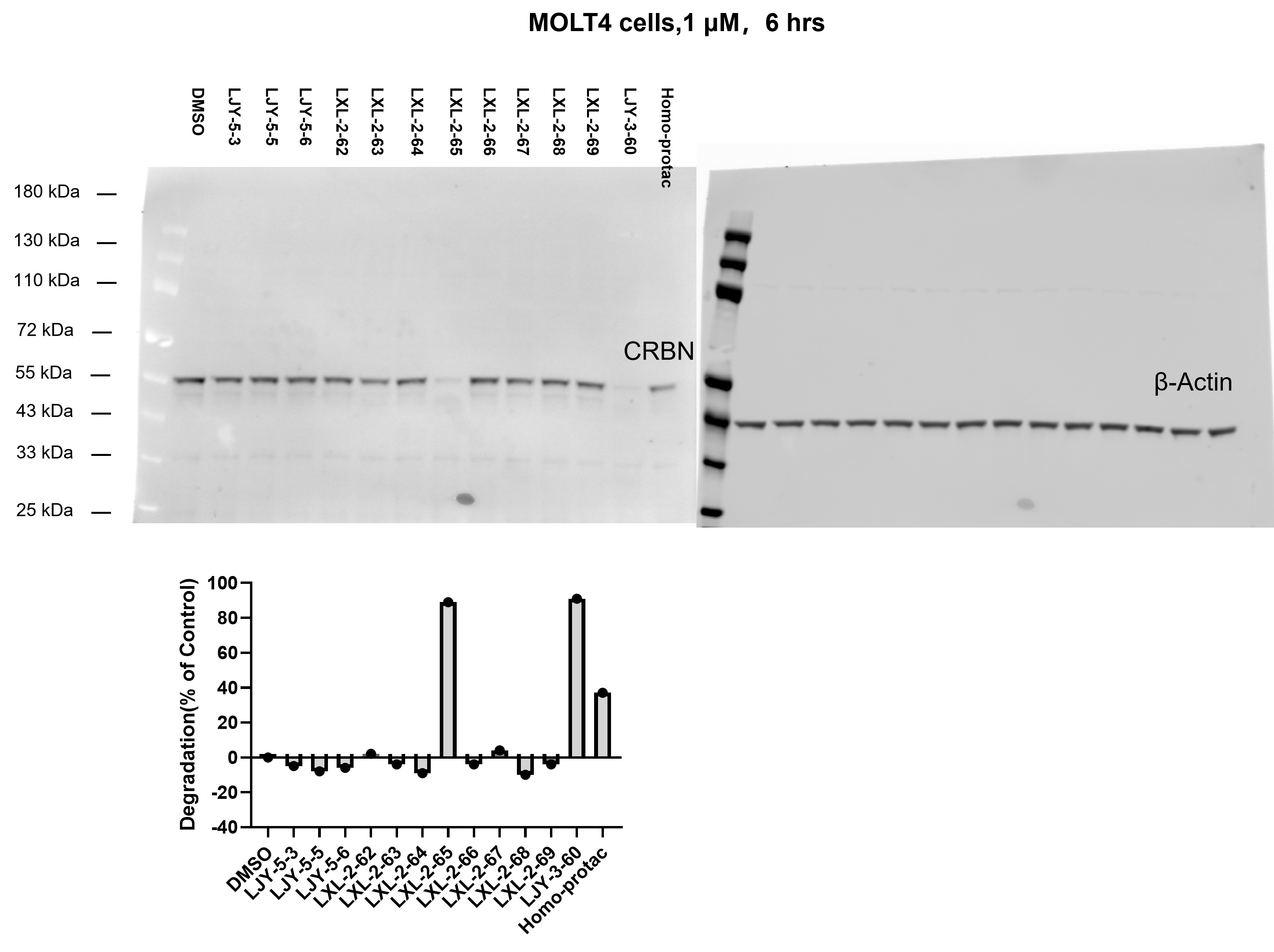

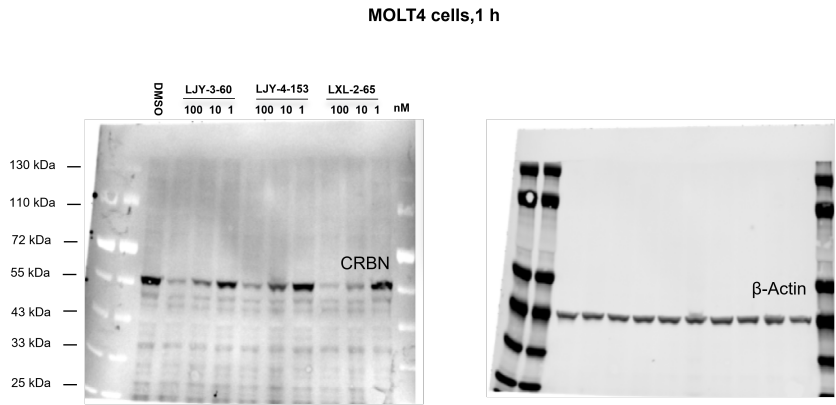

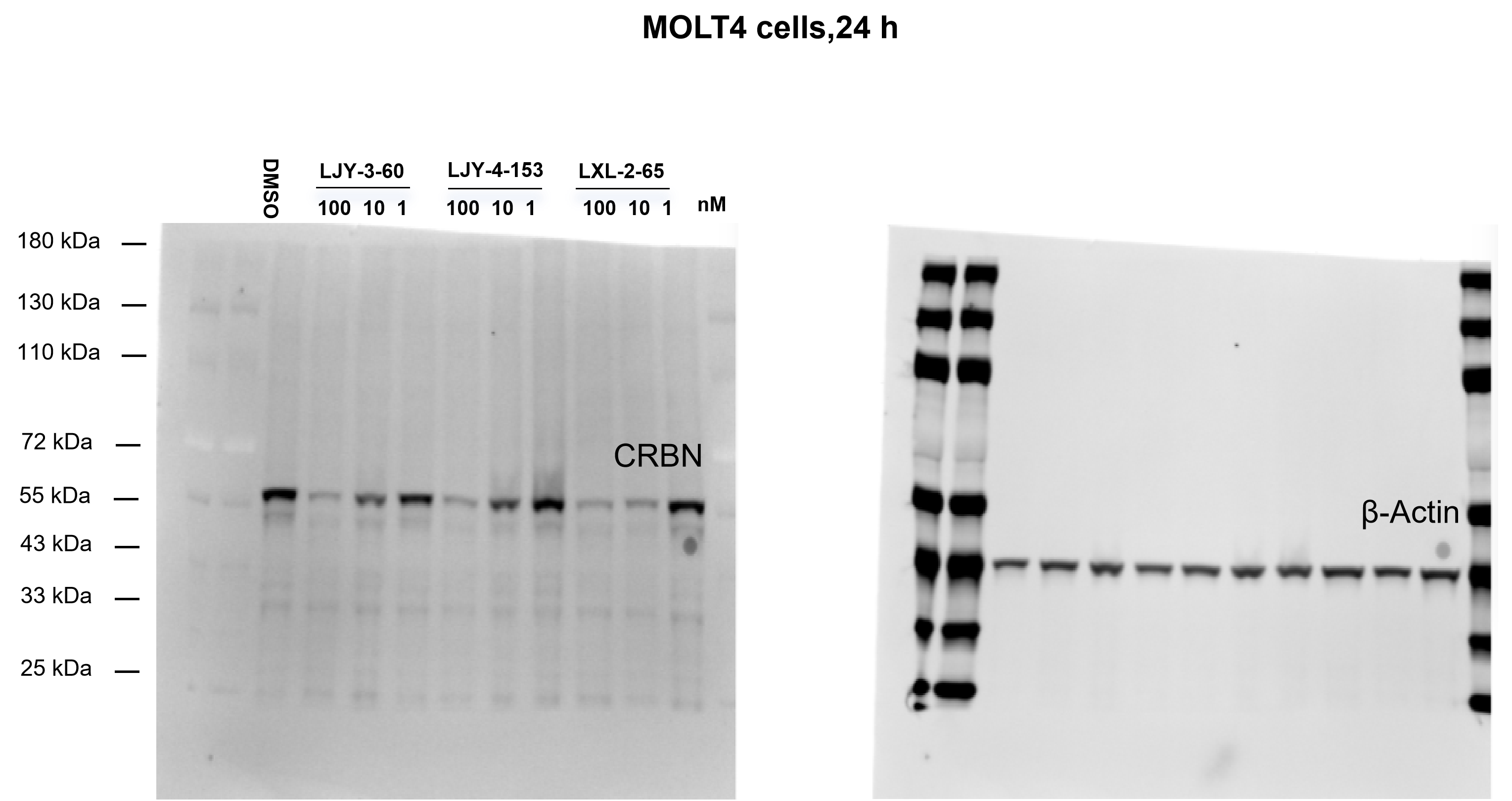

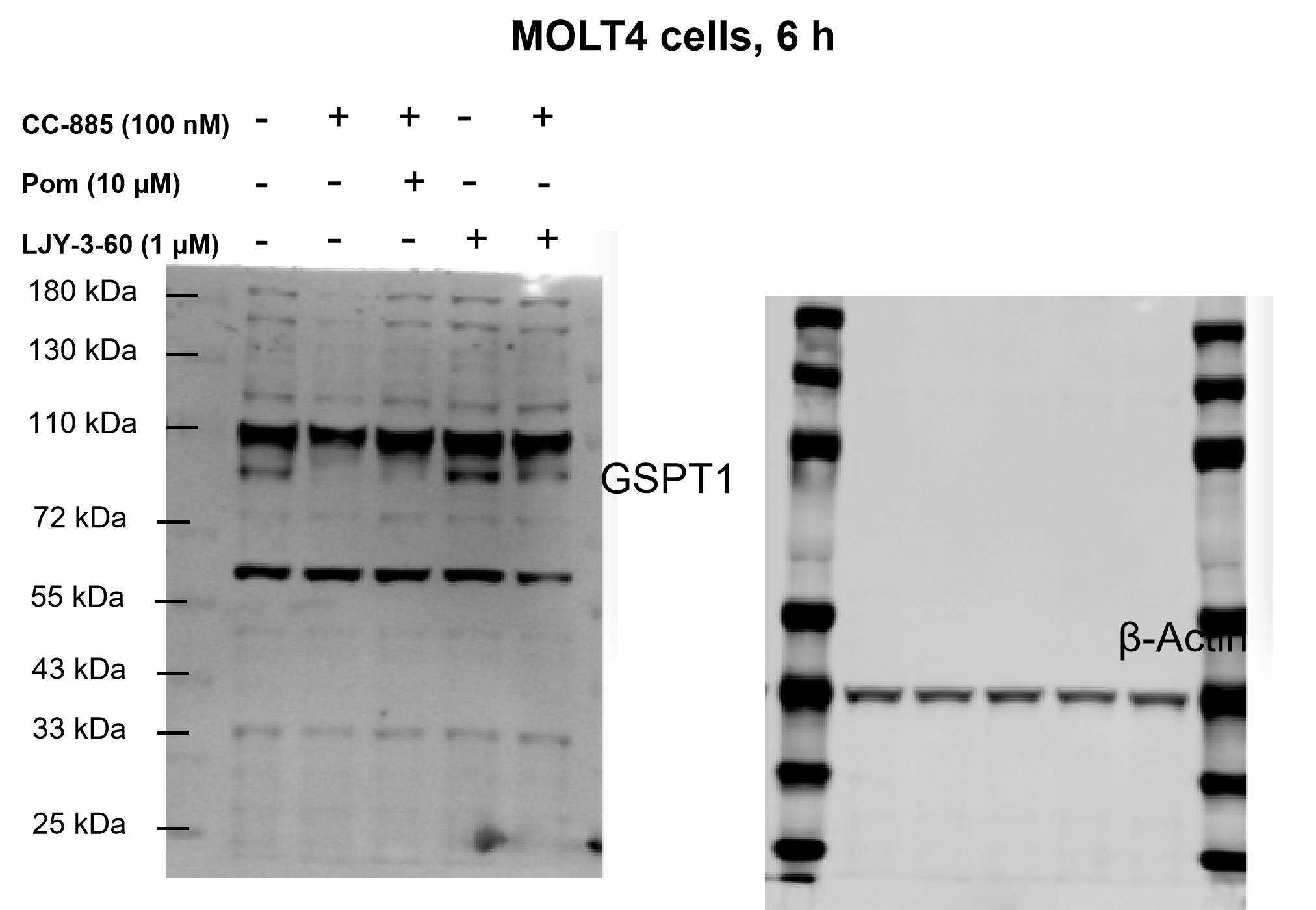

### 2. Table **S1.** Data collection and refinement statistics for crystal structure of CRBN^Midi^ complex with LJY-3-60

|  | CRBN^Midi^-LJY-3-60 (PDB: 22UL) |
| --- | --- |
| **Data Collection** |  |
| Wavelength (Å) | 0.979 |
| Resolution range (Å) | 40.0 - 1.80  (1.87 - 1.80) |
| Space group | C 1 2 1 |
| Unit cell parameters  a, b, c (Å)  α, β, γ (°) | 198.9 42.7 80.9  90.0 105.0 90.0 |
| Unique reflections | 58899 (5744) |
| Multiplicity | 5.39 (3.89) |
| Completeness (%) | 99.7 (97.9) |
| Mean I/σ(I) | 15.0 (2.10) |
| *R*_merge_ | 0.058 (0.540) |
| CC_1/2_ | 0.999 (0.875) |
| **Refinement** |  |
| *R*_work_ | 0.189 (0.296) |
| *R*_free_ | 0.222 (0.322) |
| Number of non-hydrogen atoms | 5315 |
| Macromolecules | 4972 |
| Ligands | 76 |
| Water | 224 |
| Protein residues | 635 |
| RMS bonds (Å) | 0.008 |
| RMS angles (°) | 0.860 |
| Ramachandran plot |  |
| Favoured regions (%) | 96.96 |
| Allowed regions (%) | 3.04 |
| Outliers (%) | 0.00 |
| Average B-factors (Å^2^) | 37.48 |
| Macromolecules | 37.41 |
| Ligands | 36.64 |
| Water | 39.55 |
